# A transcriptomic analysis reveals shared and inducer-specific expression patterns of cellular senescence

**DOI:** 10.64898/2026.04.20.719721

**Authors:** Jacob E. Bridge, Chen Zheng, Paul D. Robbins, Xiao Dong, Lei Zhang

## Abstract

Cellular senescence is a heterogeneous cell state induced by diverse stressors, including telomere attrition, genotoxic agents, oxidative damage, and inflammation. Despite ongoing efforts to identify conserved senescence biomarkers, it remains unclear whether senescence-inducing stimuli converge at the level of individual genes or broader molecular processes. Here, we profiled transcriptomic changes in human primary lung fibroblasts (IMR-90) driven toward senescence by replicative exhaustion, bleomycin, H_2_O_2_, or ionizing radiation under matched, dose- or time-resolved conditions. Across all four senescent inducers, global transcriptomic variation aligned along a shared axis of senescence progression, consistent with established machine learning-based senescence classifiers. However, overlap at the level of individual genes was limited, with most responses being inducer-specific or only partially conserved. In contrast, pathway-level analysis revealed far more consistent enrichment across all conditions, including downregulation of proliferation-associated pathways and activation of stress-related and pro-inflammatory pathways, accompanied by distinct inducer-specific patterns. These results support a hierarchical organization of the senescent transcriptome, in which diverse senescence inducers converge on shared pathway-level features while maintaining gene-level heterogeneity. These results provide a foundational basis for interpreting senescence signatures and may facilitate the development of more robust transcriptome-based markers of cellular senescence in aging and disease.

## Introduction

Cellular senescence is a biological program that restricts the proliferation of damaged cells (McHugh & Gil, 2018). Various stimuli can induce senescence, including telomere attrition, excessive DNA damage, oncogene overexpression, and mitochondrion dysfunction (Di Micco, Krizhanovsky, Baker, & d’Adda di Fagagna, 2021; Wiley et al., 2016). Senescence provides a crucial barrier to tumorigenesis and has been shown to facilitate development, tissue repair, and wound healing (Muñoz-Espín & Serrano, 2014). However, the accumulation of senescent cells with age has been linked to many aging phenotypes, including persistent inflammation, stem cell exhaustion, and pro-tumor microenvironments (López-Otín, Blasco, Partridge, Serrano, & Kroemer, 2023; B. Wang, Han, Elisseeff, & Demaria, 2024). Selective elimination of senescent cells can extend healthspan and lifespan in mouse models (Xu et al., 2018; Yousefzadeh et al., 2018; Zhu et al., 2015), implicating senescence as crucial factor in longevity.

Nevertheless, defining universal biomarkers of cellular senescence remains challenging. Individual molecular markers, such as p16^INK4a^, p21^Cip1^, and senescence-associated β-galactosidase (SA-β-gal), are commonly used to detect senescence, but are not present in all systems (Gorgoulis et al., 2019). Transcriptome-wide studies have identified characteristic gene expression changes associated with senescence, such as the upregulation of stress response genes, downregulation of pro-mitotic pathways, and the transition to a senescence-associated secretory phenotype (SASP) (Casella et al., 2019; Kandhaya-Pillai et al., 2023; Purcell, Kruger, & Tainsky, 2014; Wechter et al., 2023). However, these datasets also reveal a broad diversity in the molecular features of senescence, which vary by cell type, induction agent, and duration of proliferative arrest (Hernandez-Segura et al., 2017). Recent advancements, including machine learning-based classifiers trained on RNA sequencing (RNA-seq) data, have reinforced the view that senescence is best defined not by a single canonical marker but by coordinated molecular programs and signatures (Mahmud et al., 2024; Mahmud et al., 2025; Tao, Yu, & Han, 2024; J. Wang et al., 2025).

Despite this, senescence markers are often derived from heterogeneous experimental systems and datasets compiled from the literature, which, when pooled, mask much of the variation between senescence models. This makes it difficult to isolate shared and unique molecular features across different drivers of senescence. While some well-established cellular processes are conserved across different senescence inducers, it remains unclear, within a controlled experimental framework, whether the effects of these drivers converge primarily at the level of individual genes or are restricted to the level of pathways. To address these questions, we exposed human primary lung fibroblasts to increasing doses of four well-established senescence inducers: serial passaging, bleomycin, H_2_O_2_, and ionizing radiation. We then used RNA-seq data to directly compare senescence programs within this common background, revealing shared and treatment-specific signatures at the gene, pathway, and global transcriptome levels.

## Results

### Inducing dose-dependent senescence in human fibroblasts

To generate our model, we selected four senescence inducers that have been widely used within the literature. These include telomere attrition, which can result from *in vitro* serial passaging of human fibroblasts (Harley, Futcher, & Greider, 1990); bleomycin, a chemical compound that induces DNA single- and double-strand breaks (Aoshiba, Tsuji, & Nagai, 2003); H_2_O_2_, which causes oxidative damage (Chen & Ames, 1994); and ionizing radiation (IR), which leads to widespread DNA damage, including strand breaks (Suzuki et al., 2001). To induce telomere attrition, low-passage IMR-90 fibroblasts (Population Doubling Level, PDL ≈ 17) were maintained in culture until complete growth arrest, with samples collected at PDLs ≈ 17, 62, 82, or 87. For the other three inducers, IMR-90s (PDL ≈ 21) were starved in serum-free media to promote cell-cycle synchronization, then treated with 0, 1.5, 5, 15, 50, or 100 μM bleomycin; 0, 15, 50, 150, 300, or 600 μM H_2_O_2_; or 0, 1.5, 5, or 20 Gy IR (**Materials and Methods**). After 10 days, cells were harvested, stained for SA-β-gal, and subjected to RNA-seq.

IMR-90 fibroblasts exhibited dose-dependent growth arrest in response to stress (**Figs. 1A-D**). Treatments of at least 15 μM bleomycin, 150 μM H_2_O_2_, and 20 Gy IR led to persistent declines in cell count after 10 days, while lower doses yielded reduced expansion relative to controls. Exposure to 600 μM H_2_O_2_ resulted in widespread apoptosis, leading to insufficient cell density for downstream analysis. The growth rate of serially passaged IMR-90s began to slow at PDL ≈ 75, resulting in complete mitotic arrest at PDL ≈ 87. High-dose and late-passage conditions stained positive for SA-β-gal, and exhibited altered morphology, including enlarged surface area and irregular shape, consistent with cellular senescence (**Figs. S1A-D**).

**Figure 1:**
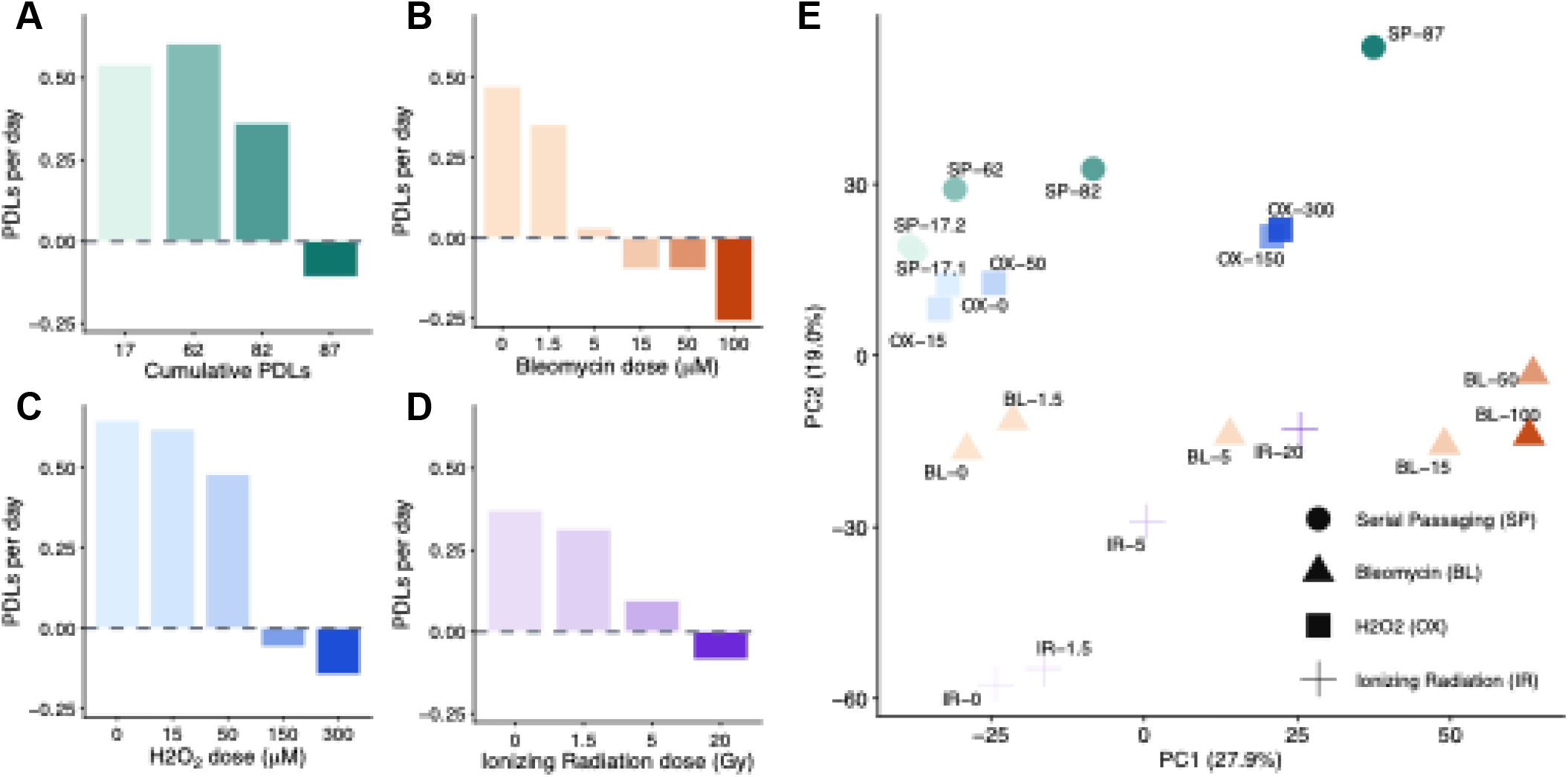
Dose-dependent senescence induction in IMR-90 fibroblasts. **(A)** Impact of serial passaging (SP), bleomycin (BL), H_2_O_2_ (OX), and ionizing radiation (IR) on growth rate. The population doubling levels (PDLs) of BL, OX, and IR samples were monitored over 10 days post-treatment. For SP, PDLs per day was calculated over the last 10 days in culture prior to harvest. **(B)** Principal component analysis (PCA) for RNA-seq data collected from SP, BL, OX, and IR samples. The first two principal components (PC1 and PC2) are displayed on the x- and y-axes, respectively. Samples are labeled by treatment and dose or PDL, with SP-17.1 and SP-17.2 serving as technical replicates for batch correction.

### Dose-dependent shifts in the senescent transcriptome

Apart from the 600 μM H_2_O_2_ condition, we performed bulk RNA-seq on the other 19 samples listed above, using two technical replicates of the low-passage cells (PDL ≈ 17) to control for batch effects during library preparation and sequencing. Libraries were sequenced on an Illumina NovaSeq X Plus system with 150 bp paired-end reads, yielding an average depth of 30.3 million reads per sample (**Table S1**), substantially higher than the required depth for detecting gene expression alterations using RNA-seq (Liu, Zhou, & White, 2014). After quality control, alignment, and expression quantification (**Materials and Methods**), we detected 32,295 genes expressed in at least one sample.

We first used principal component analysis (PCA) to examine whether transcriptome-wide expression patterns reflected a common trajectory toward cellular senescence across the four treatment types (**Fig. 1E**). The first two principal components (PCs) explained 46.9% of the total variance, with 27.9% attributed to PC1 and 19.0% to PC2 (**Fig. S2A**). High-dose conditions exhibited larger PC1 values, which displayed a strong negative correlation with growth rate across all samples (R^2^ = 0.890, *P* = 4.51 × 10^−10^; **Fig. S2B**), suggesting a unified set of senescence-associated expression changes. Along the PC2 axis, samples did not collapse into a single cluster, even among controls. (**Fig. S2C**). This was not due to batch effects during library preparation and sequencing, as the two early-passage replicates displayed similar PC1 and PC2 values, but may have partially reflected minor technical variation introduced during cell culture. However, although it did not correlate with growth rate across all samples (R^2^ = 0.0048, *P* = 0.772), PC2 strongly correlated with growth rate within non-bleomycin treatment groups (passaging: R^2^ = 0.856, *P* = 2.43 × 10^−2^; H_2_O_2_: R^2^ = 0.921, *P* = 9.66 × 10^−3^; IR: R^2^ = 0.994, *P* = 2.97 × 10^−3^; bleomycin: R^2^ = 0.066, *P* = 0.623), suggesting treatment-specific effects of senescence induction on the global transcriptome.

To determine whether global gene expression patterns observed in our dataset are consistent with existing literature, we used SenFinder, a support vector machine (SVM) classifier trained on published human fibroblast RNA-seq datasets (Mahmud et al., 2025), to infer senescence status from our data. When applied to our samples, SenFinder classified treatments of at least 5 μM bleomycin, 150 μM H_2_O_2_, 5 Gy IR, and 82 PDLs as senescent (**Fig. 2A**). PC1 correlated strongly with SenFinder senescence probability (SP) across all samples (R^2^ = 0.787, *P* = 1.87 × 10^−7^; **Fig. S3A**), suggesting that senescence-associated expression patterns shared across our treatment groups are also conserved within the literature. As with growth rate, SP did not correlate with PC2 across all samples (R^2^ = 0.049, *P* = 0.350; **Fig. S3B**), but, with the exception of bleomycin, was associated with dose-dependent variation within treatments (RS: R^2^ = 0.687, *P* = 8.30 × 10^−2^; H_2_O_2_: R^2^ = 0.908, *P* = 1.22 × 10^−2^; IR: R^2^ = 0.973, *P* = 1.38 × 10^−2^; bleomycin: R^2^ = 0.093, *P* = 0.558).

**Figure 2:**
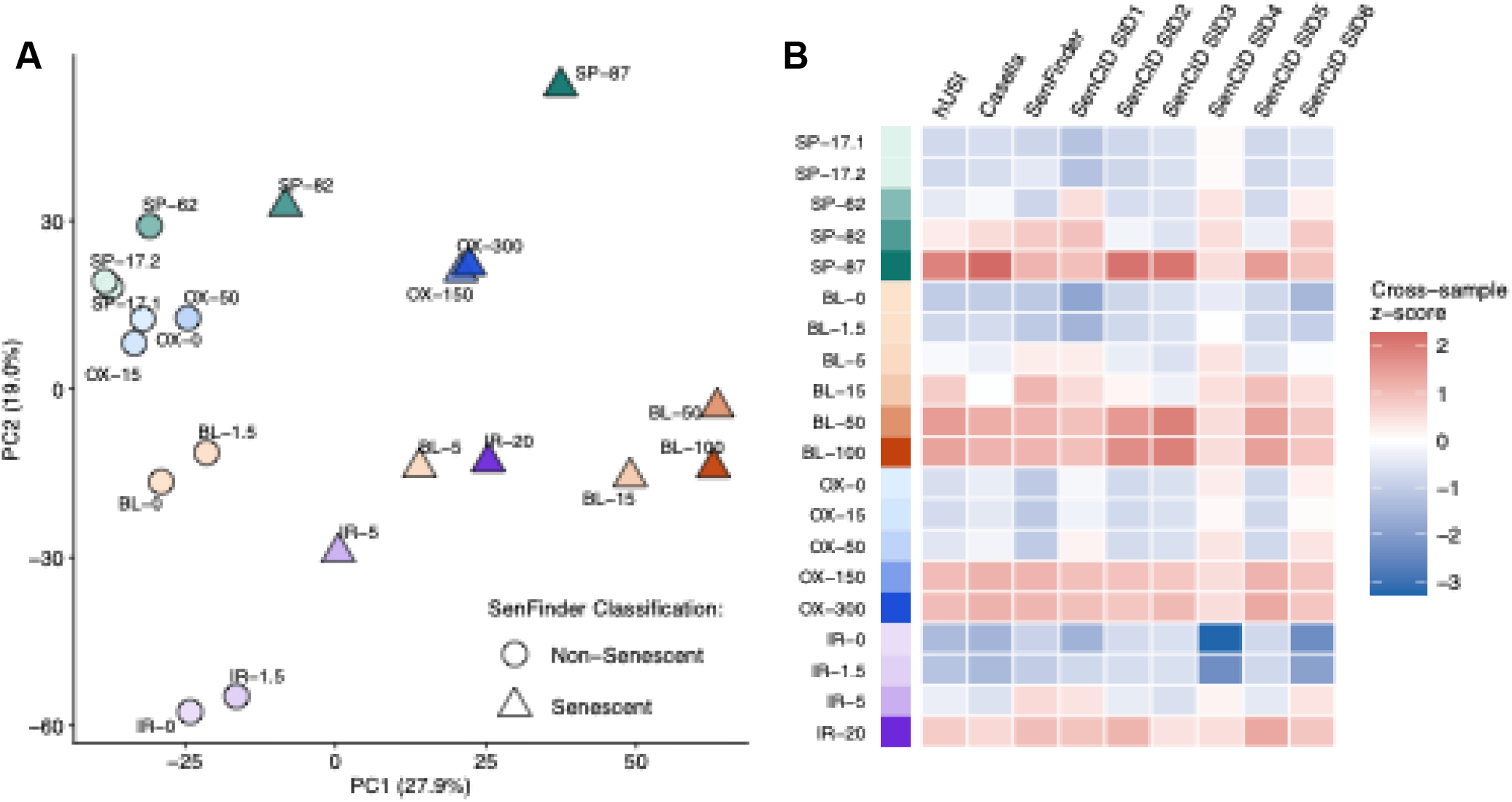
Machine learning-based classifiers detect global senescence signatures across inducers. **(A)** PCA for SP, BL, OX, and IR samples overlaid with SenFinder classification status. **(B)** Senescence classification scores generated using the human universal senescence index (hUSI), gene sets described by (Casella et al. 2019), SenFinder, or SenCID senescence identity (SID) groups. For each classifier, senescence scores were z-score-normalized across all samples.

To corroborate our findings, we applied SenCID (Tao et al., 2024), another SVM classifier, to our samples. We also computed senescence scores based on published gene sets from the human universal senescence index (hUSI) (J. Wang et al., 2025) and Casella et al. (Casella et al., 2019) (**Materials and Methods**). For SenCID, which outlines six distinct senescence identity (SID) groups, each SID was evaluated separately. Across all four scoring methods, including SenFinder, high-dose samples consistently received larger senescence scores, suggesting broad agreement between models (**Figs. 2B & S4**). Among SenCID SIDs, SID5, which has been linked to lung fibroblast senescence (Tao et al., 2024), was most strongly enriched in our IMR-90 background.

SenFinder and SID5 scores correlated most strongly with growth rate across samples (**Fig. S5**), while SID5, SID2, hUSI, and SenFinder scores correlated strongly with PC1 (**Fig. S6**). Intermediate doses, such as 15 μM bleomycin, 5 Gy IR, and 82 PDLs, produced more variable scores than other treatments, revealing the additional challenge of classifying partially senescent conditions. Models that correlated more weakly with PC1, such as the SID4, SID6, and Casella et al. feature sets, exhibited stronger correlations with PC2 (**Fig. S7**), reinforcing that, beyond culture-associated batch effects, PC2 also reflects treatment-specific enrichment of published senescence signatures. Taken together, these results indicate that senescence induction drives conserved variation in global gene expression, supplemented by stress-specific programs that differ by treatment type.

### Gene-level analysis reveals treatment-specific heterogeneity following senescence induction

After assessing global transcriptomic variation, we next examined senescence-associated expression changes within single genes. To focus on robust changes, we restricted our analysis to 16,565 genes with DESeq2-normalized counts ≥ 10 in at least three samples (Love, Huber, & Anders, 2014). To quantify differential expression within treatment types, we filtered genes based on three separate metrics: consistent directional change with dose, reflected by Spearman’s ρ; substantial upregulation or downregulation between the highest- and lowest-dose samples; and a significant linear relationship between dose and expression level. At thresholds of |ρ| ≥ 0.8, fold expression change ≥ 4, and *P* ≤ 0.05, 1,805 genes were differentially expressed in at least one treatment group (**Fig. S8A-D; Table S2**). Of these, 286 genes were differentially expressed in two treatments, 40 were differentially expressed in three treatments, and only two, *GPR68* and *SLC4A11*, were differentially expressed in all four treatments (**Fig. 3A-B**). Upregulation of *GPR68*, which promotes the expression of pro-inflammatory factors like TNF-α, was implicated as a driver of senescence in a recent review (Wei et al., 2025). Conversely, upregulation of *SLC4A11*, a sodium-coupled borate cotransporter, has not been linked to senescence in the literature, although its depletion can promote senescence in tumor cells (Y. Wang et al., 2024). Limited overlap at the gene level was consistent across all inducers, with no pair of treatments sharing more than 13% of their combined differentially expressed genes (**Fig. 3C**). Nevertheless, among differentially expressed genes shared by ≥ 2 treatments, 309 exhibited consistent directional enrichment, while only 19 displayed mixed effects, aligning with the broad similarity in global expression patterns observed between senescence inducers (**Fig. 3D**).

**Figure 3:**
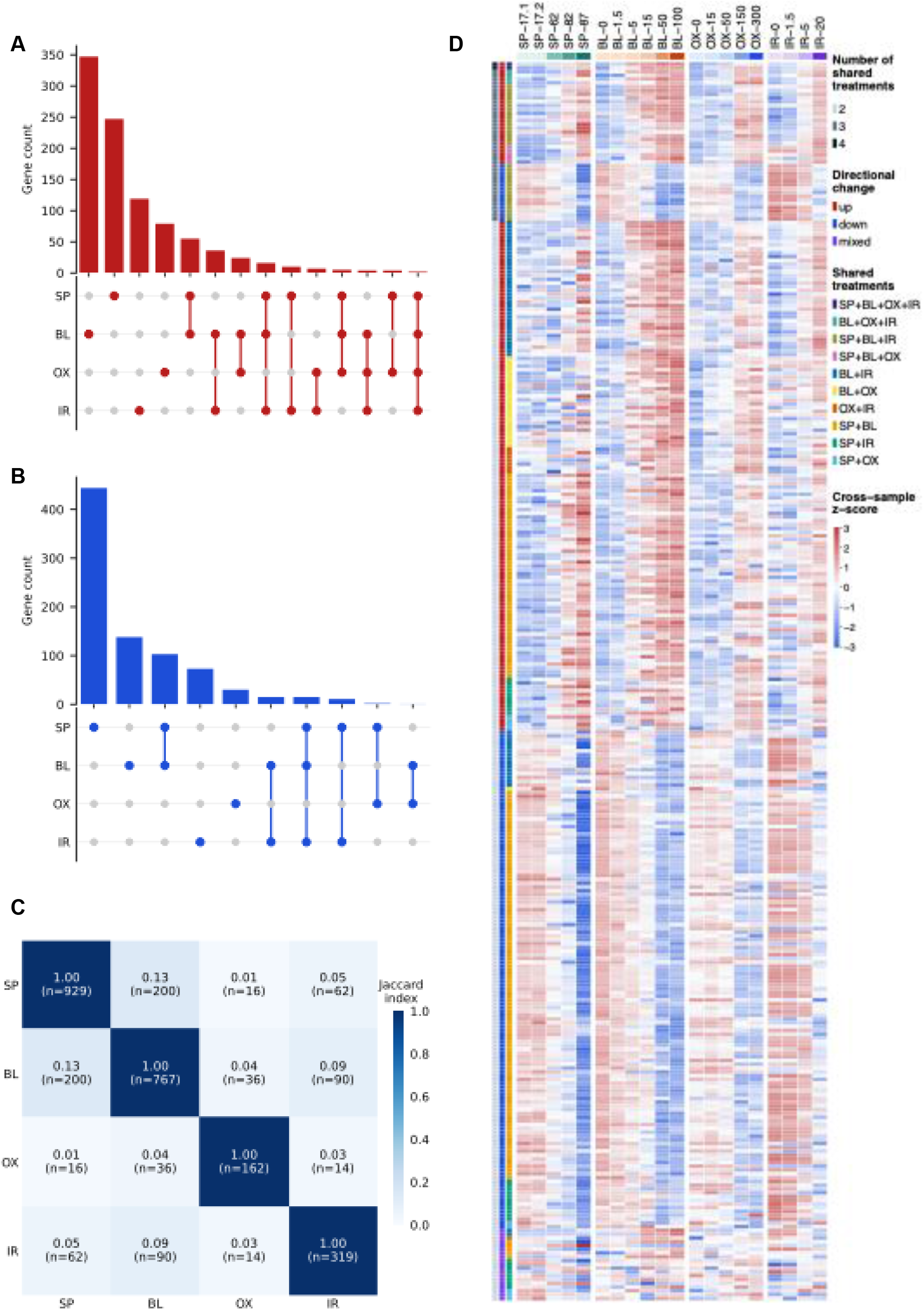
Differential gene expression is rarely shared across senescence induces. **(A)** Distribution of significantly upregulated genes induced by SP, BL, OX, and IR. **(B)** Distribution of significantly downregulated genes induced by each treatment. **(C)** Overlap of differentially expressed genes between pairs of inducers, expressed as their Jaccard index (i.e., the ratio of the number of overlaps to the number of the union). **(D)** Sample-specific expression levels of differentially expressed genes shared by more than one inducer. Genes are grouped by their combination of shared treatments, and whether they exhibit consistently increased, consistently decreased, or mixed expression changes in response to each treatment. For each gene, expression levels are z-score-normalized across all samples.

After discarding 10 entries lacking valid gene IDs, we assessed whether the remaining 299 shared, directionally consistent genes were reported in other senescence gene sets, including AgeAtlas (A. A. Consortium, 2021), CellAge (Tacutu et al., 2018), CSGene (Zhao, Chen, & Qu, 2016), HCSGD (Dong et al., 2017), and SenMayo (Saul et al., 2022). We found that genes from the current study displayed limited overlap with published gene sets (**Figs. S9A-B**). This was unsurprising given that published gene sets were also largely distinct from each other, with no two resources sharing more than 25% of their combined genes. Overall, these findings underscore one of the most pressing challenges facing the senescence field: that expression changes at the gene level are insufficient to confirm senescence across all models.

### Pathway-level analysis reveals a shared senescence program alongside inducer-specific modules

Beyond the expression changes of single genes, certain pathways, responsible for genome maintenance, cell cycle arrest, or intercellular communication, are commonly associated with senescence (Di Micco et al., 2021; Rodier et al., 2009). To determine whether enrichment of these pathways was conserved across different treatments, we performed fast gene set enrichment analysis (FGSEA) (Korotkevich et al., 2021) on our dataset. For each treatment, we ranked genes based on their Spearman’s ρ and fold-expression change, then determined enrichment of the Molecular Signatures Database (MSigDB) Hallmark (Liberzon et al., 2015), Reactome (Milacic et al., 2024), and Gene Ontology (GO) Biological Process (BP) gene sets (G. O. Consortium, 2026). Compared to 18% of single genes, 49% of differentially enriched gene sets (522/1071) were shared by at least two treatments (**Figs. 4A-B; Table S3**), including 158 enriched in three treatments and 120 enriched in all four treatments. This trend was particularly pronounced in negatively enriched gene sets, of which 58% (304/528) were enriched in at least two treatments and 20% (106/528) were enriched in all four treatments. Unlike single genes, none of the pathways exhibited significant mixed behavior across treatments.

**Figure 4:**
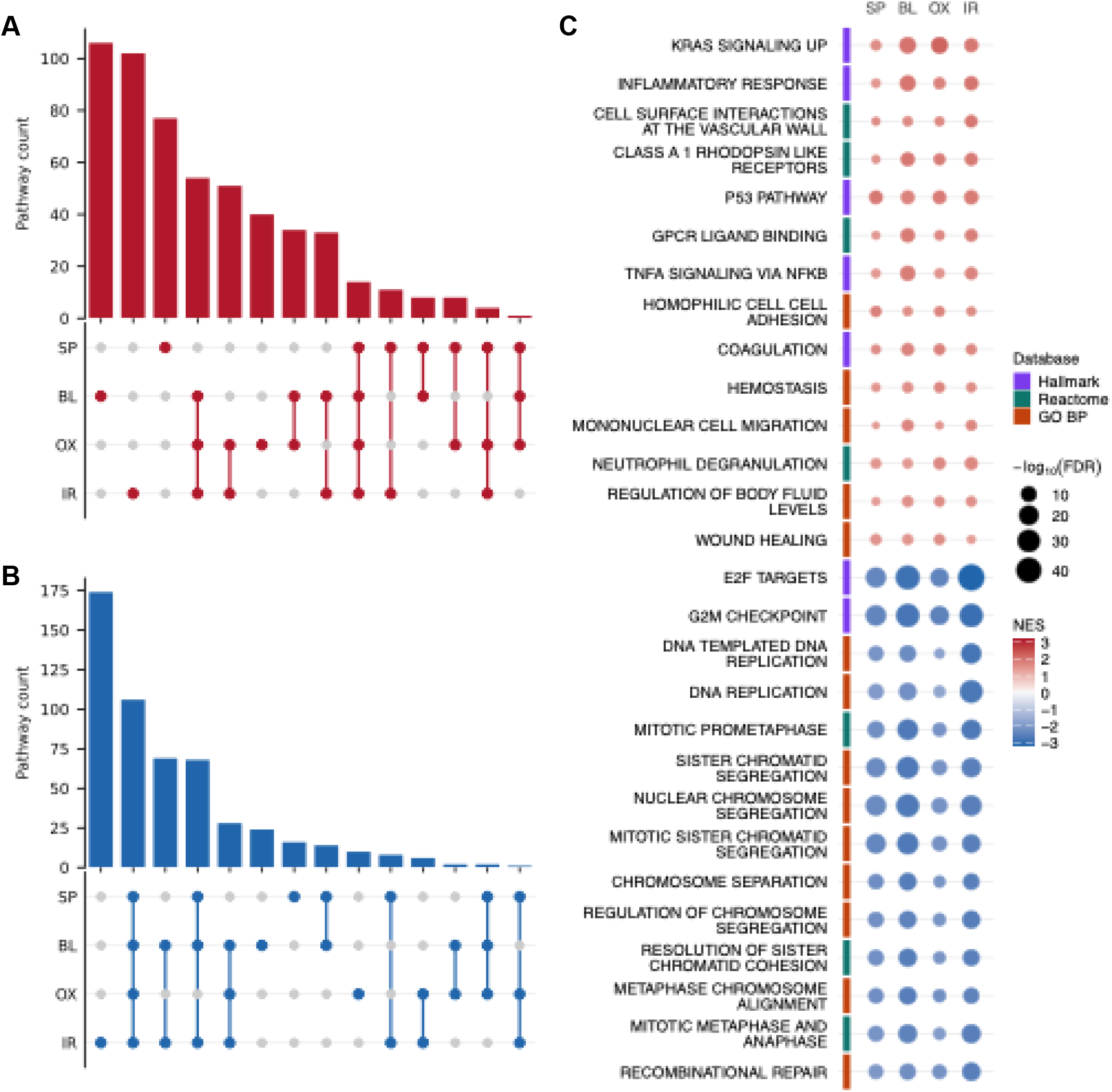
Senescence inducers are united by shared pathway enrichment. **(A)** Distribution of positively enriched gene sets induced by SP, BL, OX, and IR, determined by Fast Gene Set Enrichment Analysis (FGSEA). **(B)** Distribution of negatively enriched gene sets induced by each treatment. **(C)** Top 14 positively and negatively enriched gene sets shared across all four treatments. Pathways were ranked by their average treatment-specific normalized enrichment scores (NESs), with treatment-specific significance thresholds of Benjamini-Hochberg-adjusted False Discovery Rate (FDR) ≤ 0.05. Gene sets are labeled with their names and databases of origin.

Gene sets impacted by all four treatments aligned closely with senescence phenotypes described in the literature (**Fig. 4C**). DNA damage response elements, such as the Hallmark “P53 Pathway,” were positively enriched by all treatments. Correspondingly, gene sets associated with cell cycle progression, including “E2F Targets,” “DNA Replication,” and “G2M Checkpoint,” were negatively enriched by all treatments, mirroring the well-established link between DNA damage sensing and growth arrest observed during senescence (Di Micco et al., 2021; Kandhaya-Pillai et al., 2023). A variety of pro-inflammatory gene sets, such as “TNF-α Signaling via NF-KB” and “Neutrophil Degranulation,” were also positively enriched in all treatments, suggesting that all four senescent inducers were united by robust SASP induction. We also examined gene sets that were only enriched in one of the four treatments (**Fig. S10A**). Despite a few consistent trends, such as the negative enrichment of ribosome-associated gene sets following bleomycin treatment, stress-specific enrichment patterns appeared largely sporadic, spanning a wide variety of cellular functions.

To more selectively identify pathways exhibiting treatment-dependent variation, we ranked Hallmark gene sets based on the standard deviation of their normalized enrichment scores (NES) across treatments. We restricted our analysis to gene sets with Benjamini-Hochberg-adjusted False Discovery Rate (FDR) ≤ 0.05 in at least one treatment, and performed two separate analyses: one using raw NES, and another in which NES was set to zero for treatments with FDR > 0.05.

Among the 20 most variable gene sets produced by each ranking, 15 high-confidence entries were shared between both lists (**Fig. S10B**). Some gene sets, such as “Hypoxia,” Epithelial-to-Mesenchymal Transition (“EMT”), and “MYC Targets V2,” differed in magnitude between the inducers, but remained directionally consistent across all treatments (**Figs. 5A-D**). Conversely, several pro-inflammatory pathways, such as “IFNA Response,” “IFNG Response,” and “IL6-JAK-STAT3,” were positively enriched to varying degrees following bleomycin, H_2_O_2_, or IR treatment, but were non-significantly depleted during serial passaging. Overall, Bleomycin and IR led to similar differential enrichment of highly variable pathways, which differed marginally from the effects of H_2_O_2_, and more strongly from the impact of serial passaging. This mirrored the degree of overlap across all differentially enriched gene sets, with Bleomycin and IR producing the highest fraction of shared pathways, and serial passaging displaying the lowest average similarity to other inducers (**Figs. 5A-D**). Nevertheless, between any combination of treatments, gene sets were far more conserved than individual genes (**Fig. 5E**), suggesting that distinct drivers of cellular senescence predominantly converge at the pathway level.

**Figure 5:**
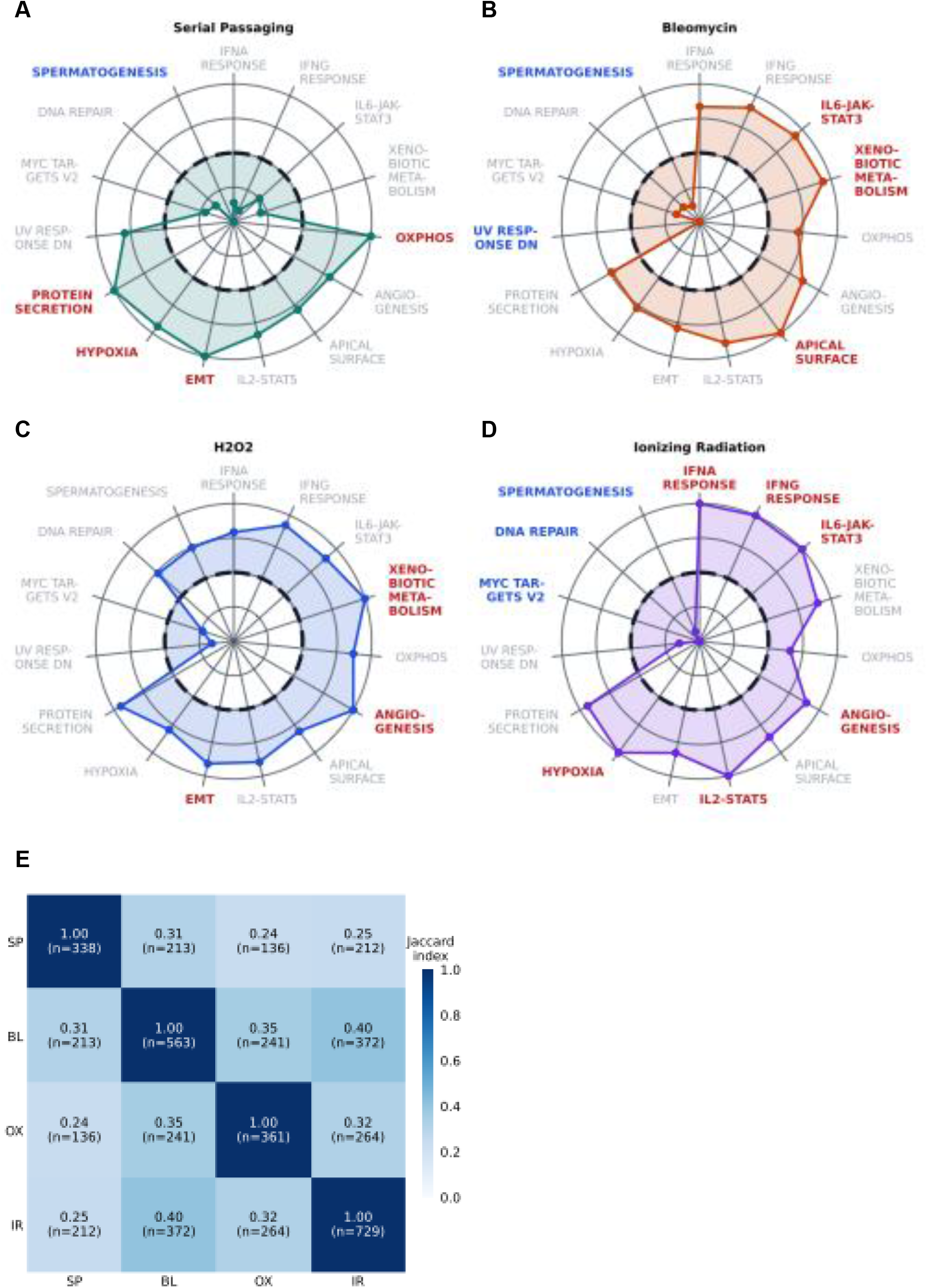
Highly variable pathways reveal inducer-specific senescence signatures, despite broad pathway-level similarity. **(A-D)** Relative enrichment of 15 highly variable Hallmark gene sets following SP, BL, OX, or IR treatment. For each gene set, inducer-specific NESs were normalized to the highest |NES| across all inducers, then plotted relative to a dashed circle denoting zero normalized enrichment. Significant positive and negative enrichment are denoted by red and blue labels, respectively, while gray labels indicate gene sets that did not achieve a Benjamini-Hochberg-adjusted FDR ≤ 0.05 for a given treatment. **(E)** Overlap of differentially enriched gene sets between pairs of inducers, expressed as their Jaccard index (i.e., the ratio of the number of overlaps to the number of the union).

## Discussion

Senescence is broadly defined by several hallmarks, such as proliferative arrest, the SASP, apoptosis resistance, and altered metabolism (Hernandez-Segura, Nehme, & Demaria, 2018). However, widespread heterogeneity across cell types and stress mechanisms has made it difficult to define shared markers of cellular senescence. For example, SA-β-gal, one of the most widely used senescence markers, is also induced by temporary quiescence, and is frequently expressed by non-senescent macrophages (Shah, Al-Hashimi, Benedetto, & Ruchaya, 2025). Recently, large-scale efforts to define core senescence factors based on transcriptomic data have gained increasing popularity, but still suffer from a variety of limitations. RNA-seq datasets compiled from the literature, such as those informing SenCID (Tao et al., 2024) and hUSI (J. Wang et al., 2025), use harmonized computational pipelines and sequencing batch correction to identify common trends across senescence models. However, these analyses cannot control for variation in culture materials, technique, or laboratory environment across different experiments. Gene panels that are manually curated from the literature, such as CellAge (Tacutu et al., 2018) and SenMayo (Saul et al., 2022), face the same issue. Thus, the limited overlap among these resources may not only reflect the biological heterogeneity of senescence, but also a lack of reproducibility that has long hindered progress in the field.

We addressed these concerns by assessing the dose-dependent effect of four senescence inducers on gene expression patterns within the same IMR-90 fibroblast model. At the global transcriptomic level, dose, which correlated negatively with growth rate, had the single greatest effect on between-sample variation, suggesting a unified set of transcriptional changes associated with senescence. However, this did not explain the majority of between-sample variation, reinforcing existing literature suggesting that different inducers produce distinct senescence phenotypes (Cohn, Gasek, Kuchel, & Xu, 2023; Wiley et al., 2017). Regardless of treatment, senescence classification by transcriptome-based machine learning models strongly correlated with growth rate, as well as the major principal components of global expression variation. In some cases, samples with a positive growth rate, such as 5 μM bleomycin or 5 Gy IR were, nevertheless, classified as senescent. In these samples, heterogenous morphological changes and SA-β-gal staining (**Fig. S1**), coupled with reduced expansion relative to controls, suggested that a fraction of the cells had become senescent, while the rest continued to proliferate. Thus, while individual senescence markers isolated from multi-study datasets may not be reproducible, machine learning models trained on these datasets can robustly detect senescence signatures in bulk samples.

Even in previous studies (Casella et al., 2019; Purcell et al., 2014) in which senescence induction methods are compared within the same experimental framework, emphasis is largely placed on the behavior of single genes. While confirming that differentially expressed genes vary widely between treatments, we showed that a far greater degree of overlap emerges at the pathway level. This discrepancy likely reflects a combination of biological and technical factors. At the statistical level, single-gene counts are less robust to stochastic variation than whole-pathway rankings, and are more sensitive to arbitrary filtering thresholds. This may explain why several well-characterized senescence genes, such as *p21*^*Cip1*^ and *SERPINB2*, were detected in only three out of four treatments. As in other studies (Casella et al., 2019), we sought to address this by employing less stringent statistical thresholds for differential expression, although this increases the likelihood of detecting false positives. At the biological level, distinct combinations of single genes, which may be uniquely sensitive to different cellular stressors, may produce similar cumulative effects by converging along the same signaling axis. Regardless of the cause, our results nonetheless highlight that shared biological pathways constitute the common core features of senescence, whereas heterogeneity is more pronounced at the level of individual genes.

Despite these findings, our study faces some limitations. We conducted experiments in a single human fibroblast background, and the extent to which gene and pathway-level variation is conserved across different cell types remains to be determined. In addition, our analysis does not establish causal relationships between shared pathways and senescence phenotypes, or attempt to explain the mechanisms behind observed stress-specific variation. Similarly, we did not test whether pathway-level variation, such as differential enrichment of inflammation-associated gene sets, led to treatment-specific changes at the protein level, which could have been monitored by assessing the production of SASP factors. Nevertheless, these results support a model in which diverse senescence induction methods converge on a shared pathway-level program while preserving stress-dependent variation. This framework may offer a more practical basis for defining and comparing senescent states than individual gene markers alone, a point reinforced by the success of transcriptome-based classifier studies.

## Materials and Methods

### Cell culture

Early-passage (PDL ≈ 12) IMR-90 human lung fibroblasts were obtained from the NIA Aging Cell Culture Repository at the Coriell Institute. IMR-90 cells were grown in Eagle’s Minimum Essential Media (EMEM; ATCC, 30-2003) containing 10% FBS, 100 U/mL penicillin, and 100 μg/mL streptomycin (1% P/S), and maintained at 37 °C with 5% CO_2_ and 3% O_2_.

### Senescence induction

To induce replicative senescence (RS), IMR-90 fibroblasts were serially passaged until sustained growth arrest, indicated by a reduction in cell count over two consecutive passages, was observed. Cell pellets were harvested after 10, 92, 138, or 193 days in culture (PDLs ≈ 17, 62, 82, or 87, respectively). For bleomycin, H_2_O_2_, and ionizing radiation (IR)-induced senescence, fibroblasts were seeded at 1.8 × 10^4^ cells/cm^2^ in complete EMEM four days before treatment. Three days later, cells were starved in serum-free media (0% FBS, 1% P/S) to induce quiescence. After 24 hours, cells were washed twice with phosphate-buffered saline, and treated, in base EMEM (0% FBS, 0% P/S), with different doses of Bleomycin (0, 1.5, 5, 15, 50, or 100 μM), H_2_O_2_ (0, 15, 50, 150, 300, or 600 μM), or IR (0, 1.5, 5, or 20 Gy). IR was supplied by an RS-2000 X-ray irradiator (Rad Source Technologies) at a rate of 1.8 Gy/min, while Bleomycin and H_2_O_2_ exposures lasted 2 hours.

Following treatment, cells were trypsinized, harvested, and re-seeded in complete EMEM at 1.8 × 10^4^ cells/cm^2^. Because a single H_2_O_2_ treatment was not sufficient to induce senescence in the early-passage IMR-90s (data not shown), H_2_O_2_ conditions received two additional treatments separated by 24-hour intervals. The second and third treatments did not involve serum starvation and were followed by a media change rather than re-seeding, but were otherwise identical to the first. Cells were maintained in culture for 10 days post-senescence induction–relative to the first treatment for H_2_O_2_ samples–during which 4,000 cells per sample were re-seeded in 24-well plates for senescence-associated β-galactosidase staining. The remaining cells were pelleted on day 10 and stored at -80 °C alongside the RS samples.

### Senescence-associated β-galactosidase (SA-β-gal) staining

After being re-seeded in 24-well plates, cells were temporarily maintained in culture (≥24 hours for proliferating conditions; 5 days for non-proliferating conditions) to ensure robust adhesion and preserve morphological characteristics. On day 10 post-senescence induction, cells were prepared as directed by a Senescence β-galactosidase Staining Kit (Cell Signaling Technology). Briefly, cells were fixed for 12 minutes at room temperature, then treated with a β-galactosidase staining solution containing 1 mg/mL X-gal. Plates were incubated for 20 hours at 37 °C under ambient atmospheric conditions, then treated with a 1:1 mixture of DMSO and H_2_O to remove crystals. Staining was visualized using a CKX53 Inverted Phase Contrast Microscope (Olympus).

### RNA isolation and sequencing

RNA was harvested from cell pellets using a Quick-DNA/RNA Microprep Plus Kit (Zymo). Total RNA was delivered to Novogene Inc., where it was subjected to mRNA library preparation with poly-A enrichment. Libraries were sequenced on the NovaSeq X Plus sequencing platform for 2 × 150-bp paired-end reads by Novogene.

### Data processing and analysis

For quality control, we subjected raw sequencing reads from each sample to FastQC (version 0.11.9) (Wingett & Andrews, 2018). Transcriptome alignment (to the reference genome: GRCh38) was performed using STAR (version 2.7.9a) (Dobin et al., 2013). Alignments were processed using Picard tools (Broad Institute), and gene expression was quantified using RSEM (version 1.3.1) (Li & Dewey, 2011). Gene counts were normalized in R using DESeq2 (version 1.46.0) (Love et al., 2014), and all downstream analysis was performed in R and Python.

### Machine learning methods to predict senescent status

For SenFinder (Mahmud et al., 2025), we applied the SenFinder classifier directly to our dataset. TPM values for the 20 required model features (genes) were used. The released senFinder.rds model was used to generate predicted classes and class probabilities.

For SenCID (Tao et al., 2024), we applied the SenCID model directly to our bulk RNA-seq data in the following steps. Gene expression values were normalized to counts per million, transformed as log2(CPM + 1), and z-scored within each sample. The processed matrix was then evaluated using the published SID1–SID6 models separately.

For hUSI (J. Wang et al., 2025), we derived a senescence score from the published weighted feature list rather than applying a separately released classifier object, which was not available from the original paper. Normalized expression values were transformed as log2(normalized count + 1). For each sample, we calculated the Spearman correlation between the sample expression values and the published hUSI feature weights across overlapping genes, and then min–max scaled the resulting values across samples to obtain the final hUSI score.

For the Casella et al (Casella et al., 2019), we used the published overexpressed and underexpressed gene sets rather than a directly released prediction model, which was not available from the original paper. Using the same gene-symbol matrix of log2(normalized count + 1) expression values, we computed a rank-based single-sample score for each gene set. The final Casella score was defined as the overexpressed score minus the underexpressed score.

## Data availability

The RNA-seq data generated in this study have been submitted to the NCBI Sequence Read Archive under submission ID SUB16119965, and accession number will be provided upon release.

## Acknowledgements

This work was supported by the U.S. National Institutes of Health (R35 GM159832, P01 HL160476, U19 AG056278, U54 AG079754, U54 AG076041, and T32 AG029796). The funders had no role in study design, data collection and analysis, decision to publish, or preparation of the manuscript.

## Notes

### Competing Interest Statement

The authors have declared no competing interest.

