## Supplemental Figs for "A transcriptomic analysis reveals shared and inducer-specific expression patterns of cellular senescence"

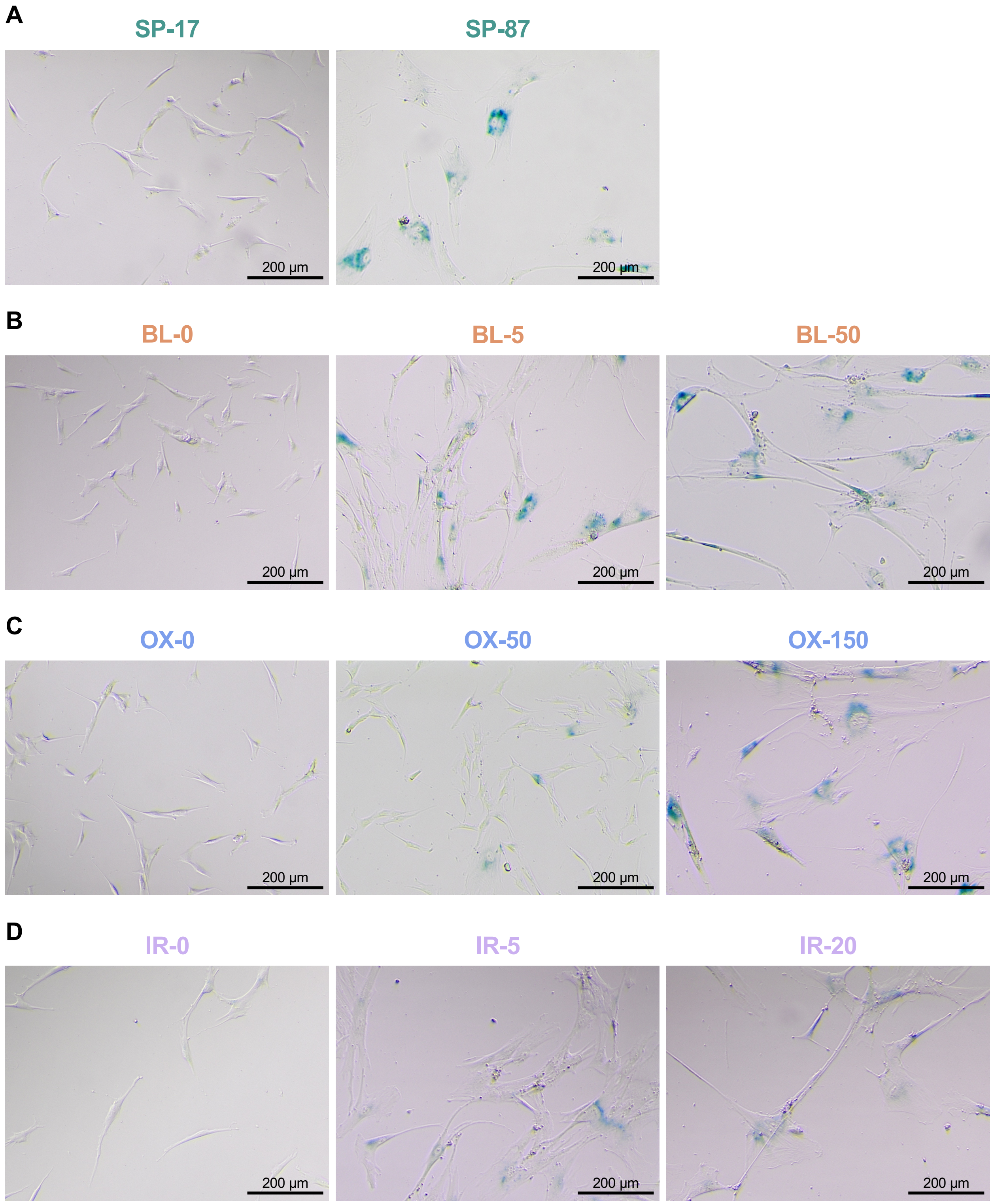


**Figure S1: Morphological changes and senescence-associated β-galactosidase (SA-β-gal) staining increase with dose and time in culture. (A-D)** IMR-90 fibroblasts subjected to serial passaging (SP), bleomycin (BL), $H_2O_2$ (OX), or ionizing radiation (IR), then stained for SA-β-gal expression. Left-to-right: BL, OX, and IR include representative images from low, intermediate, and high-dose samples, while SP includes early passage and senescent samples. Labels include treatment type and population doubling level (PDL) or dose.

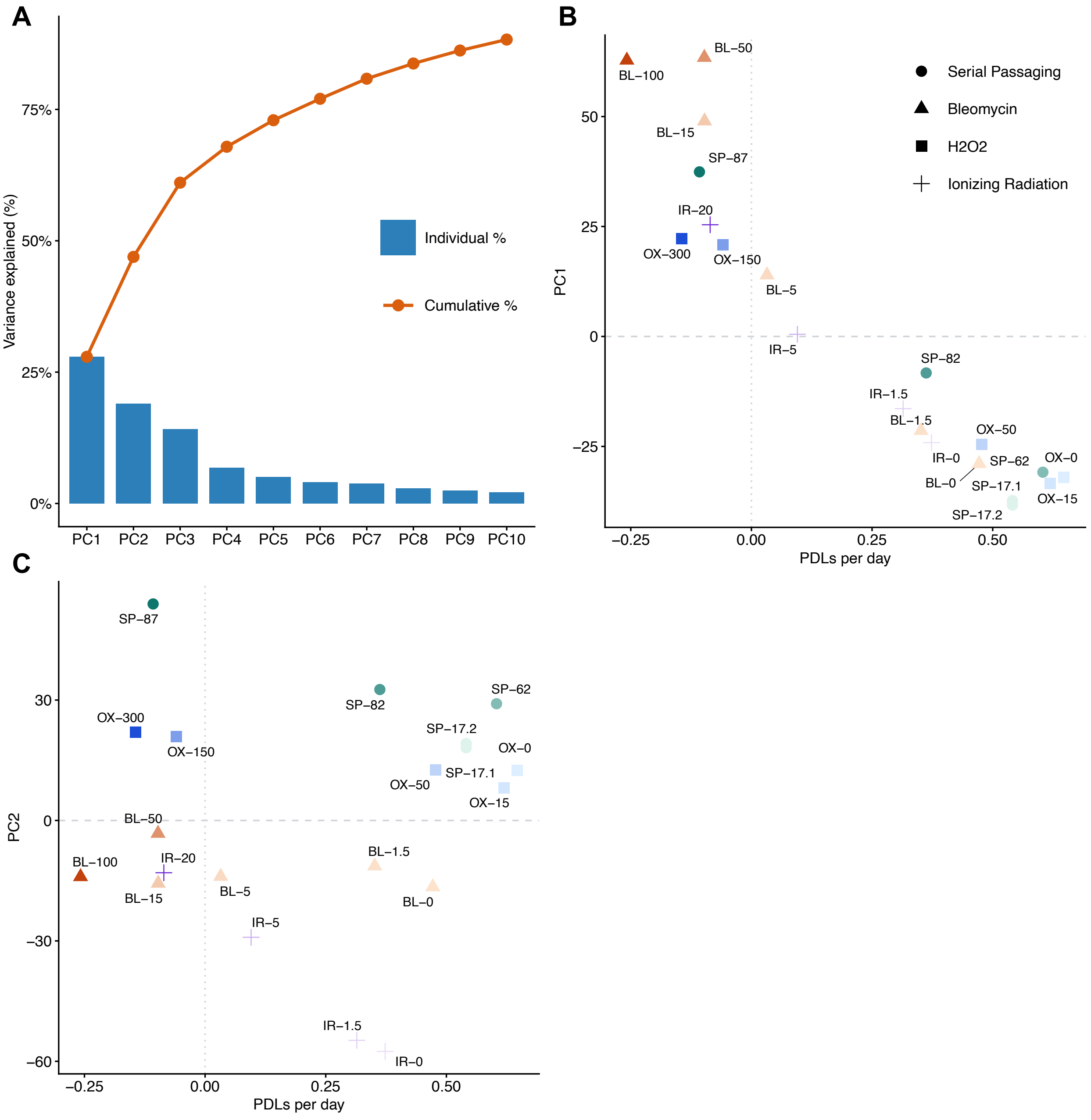


**Figure S2: Major principal components of global expression variation are associated with growth arrest. (A)** Top 10 principal components determined by a principal component analysis (PCA) of RNA-seq data from SP-, BL-, OX-, or IR-treated samples. Blue bars indicate the fraction of variance explained by each principal component, while the orange line indicates the cumulative fraction of variance up to and including the indicated component. **(B)** Principal component 1 (PC1) value as a function of growth rate for each sample. Growth rate is expressed as population doubling levels (PDLs) per day, equivalent to Fig. 1A. Samples are labeled by treatment and dose or PDL. **(C)** Principal component 2 (PC2) as a function of growth rate for each sample.

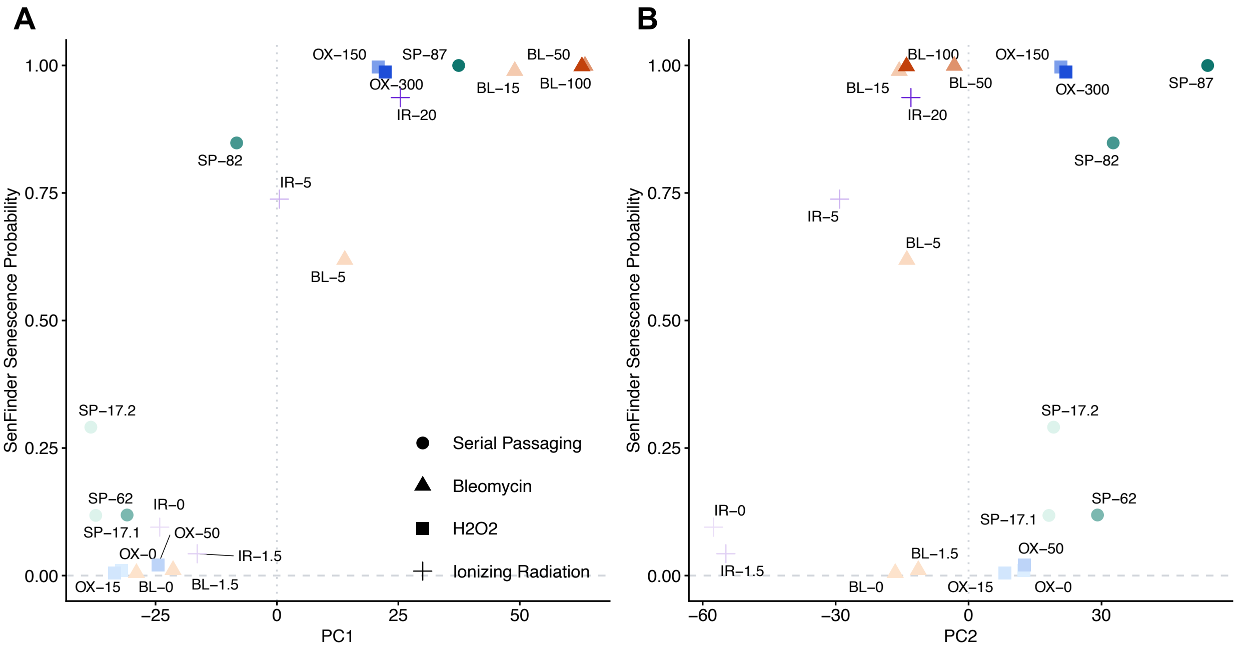


**Figure S3: Major principal components of global expression variation are associated with SenFinder senescence probability (SP). (A)** SenFinder SP as a function of PC1 (. Samples are labeled by treatment and dose or PDL. **(B)** SenFinder SP as a function of PC2.

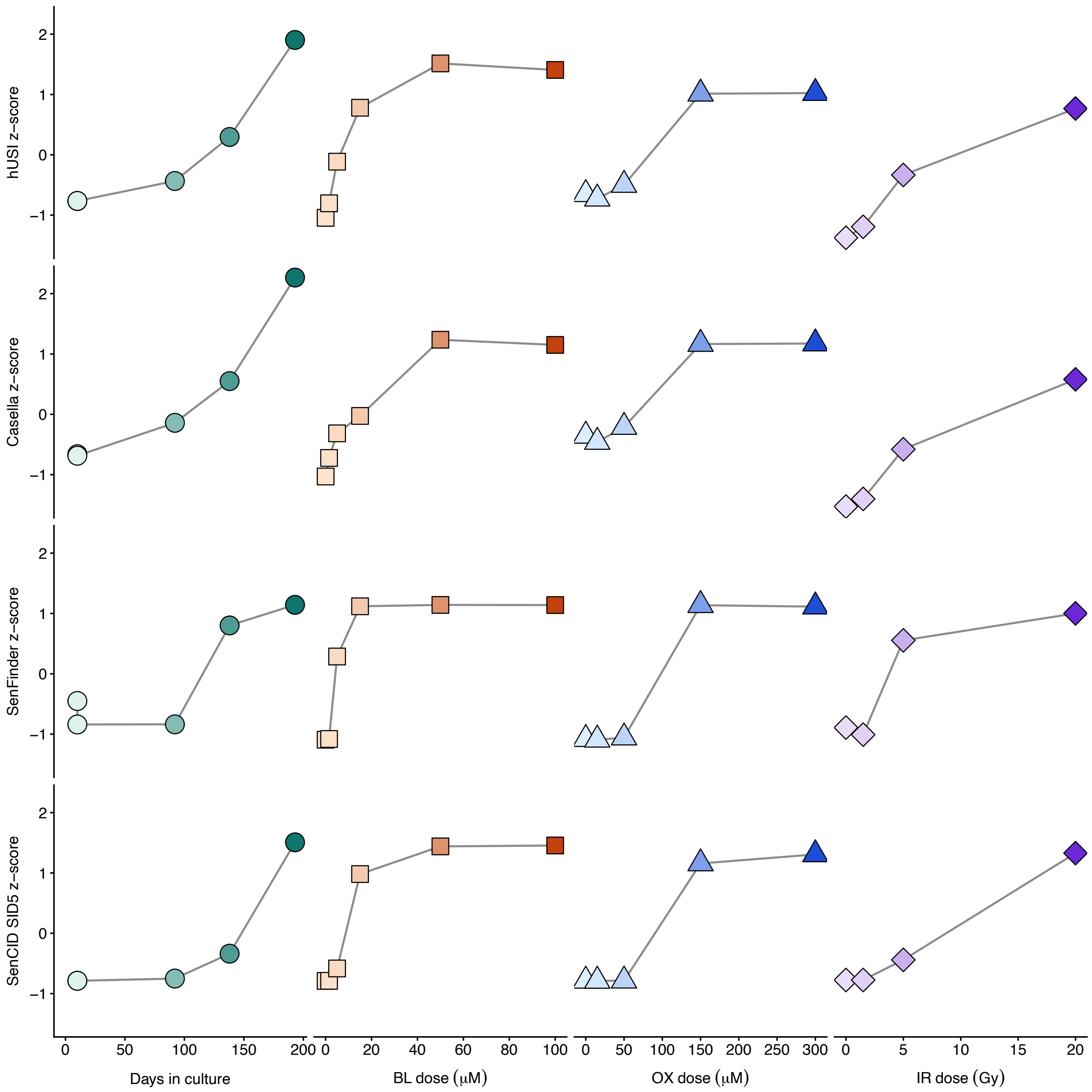


**Figure S4: Dose- and time-dependent increases in senescence score are consistent across classifiers.** Senescence classification scores were generated using the human universal senescence index (hUSI), gene sets described by Casella et al. 2019, SenFinder, or SenCID senescence identity 5 (SID5). For each classifier, senescence scores were z-score-normalized across all samples. Data points in this figure are equivalent to the cells from Fig. 2B.

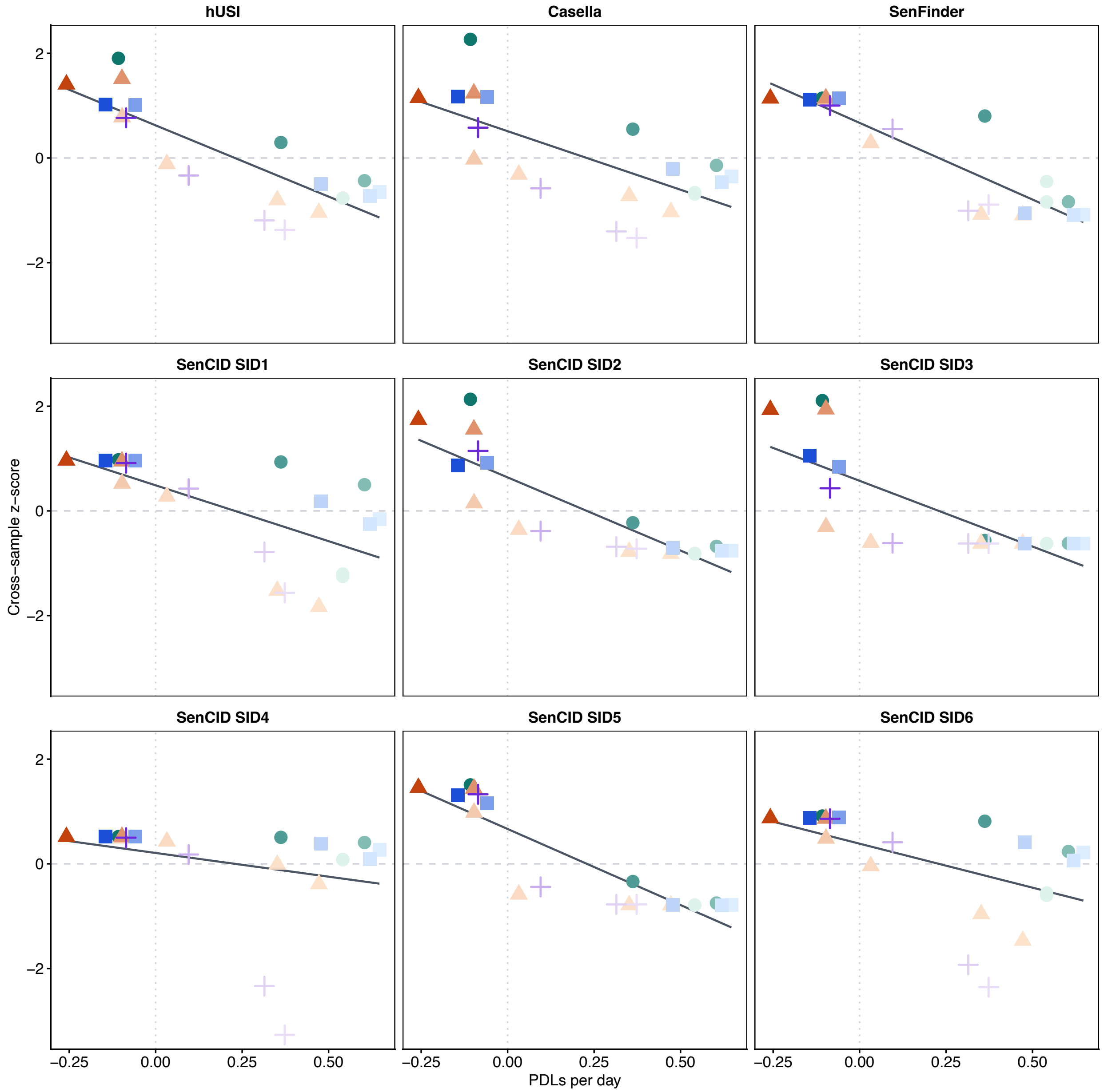


**Figure S5: Senescence classification scores are inversely correlated with growth rate.** Z-score-normalized senescence scores from hUSI, Casella et al. 2019, SenFinder, and SenCID SIDs (Fig. 2B, Fig. S4) are expressed as a function of PDLs per day (Fig. 1A). Treatment type is indicated by shape (SP: circle; BL: square; OX: triangle; IR: plus-sign) and color (SP: green; BL: orange; OX: blue; IR: purple), with darker colors corresponding to higher doses.

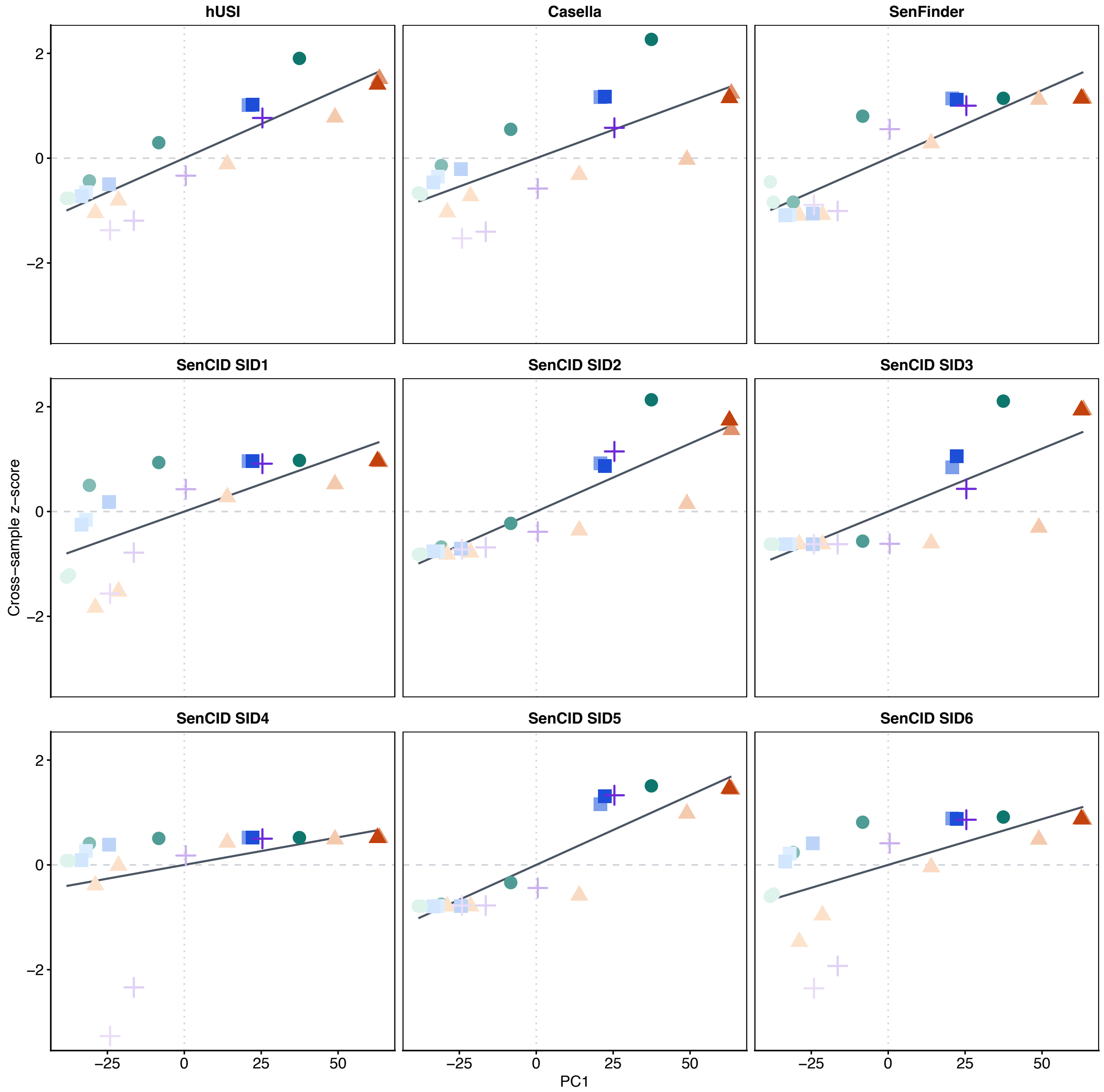


**Figure S6: Senescence classification scores correlate positively with PC1.** Z-score-normalized senescence scores from hUSI, Casella et al. 2019, SenFinder, and SenCID SIDs (Fig. 2B, Fig. S4) are expressed as a function of PC1 (Fig. 1B). Treatment type is indicated by shape (SP: circle; BL: square; OX: triangle; IR: plus-sign) and color (SP: green; BL: orange; OX: blue; IR: purple), with darker colors corresponding to higher doses.

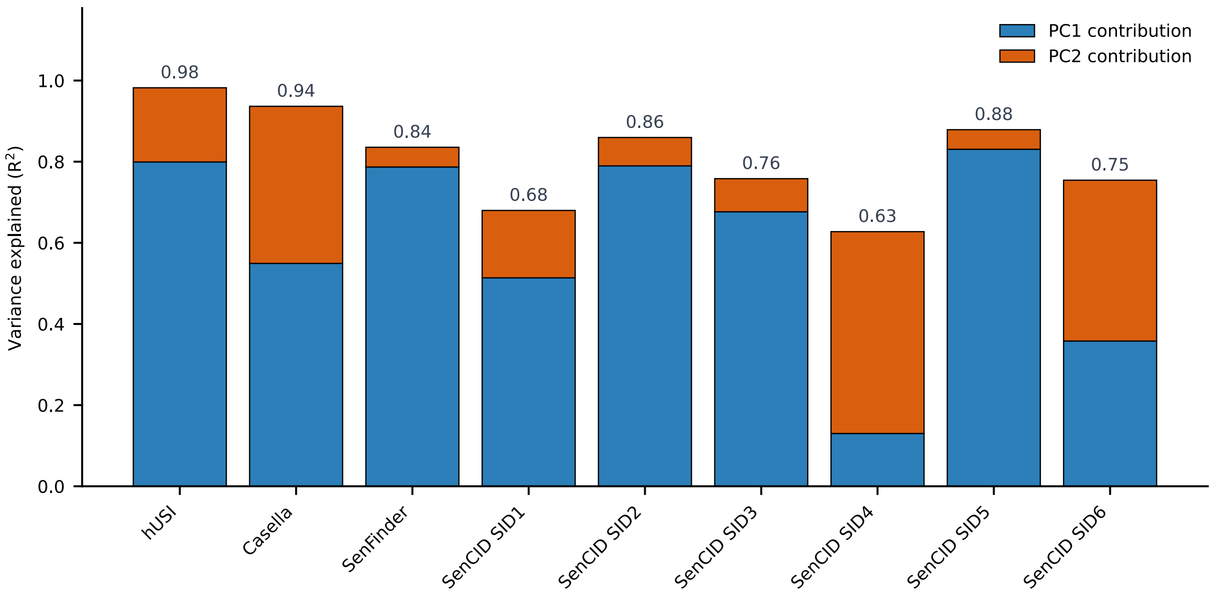


**Figure S7: Major principal components of global expression variation explain most of the variance in senescence classification scores.** Blue bars indicate the squared Pearson correlation ($R^2$) between PC1 (Fig. 1B) and senescence scores from hUSI, Casella et al. 2019, SenFinder, and SenCID SIDs (Fig. 2B, Fig. S4). Orange bars indicate $R^2$ between PC2 (Fig. 1B) and senescence scores. Bars are labeled with the sum of the variance explained by PC1 and PC2.

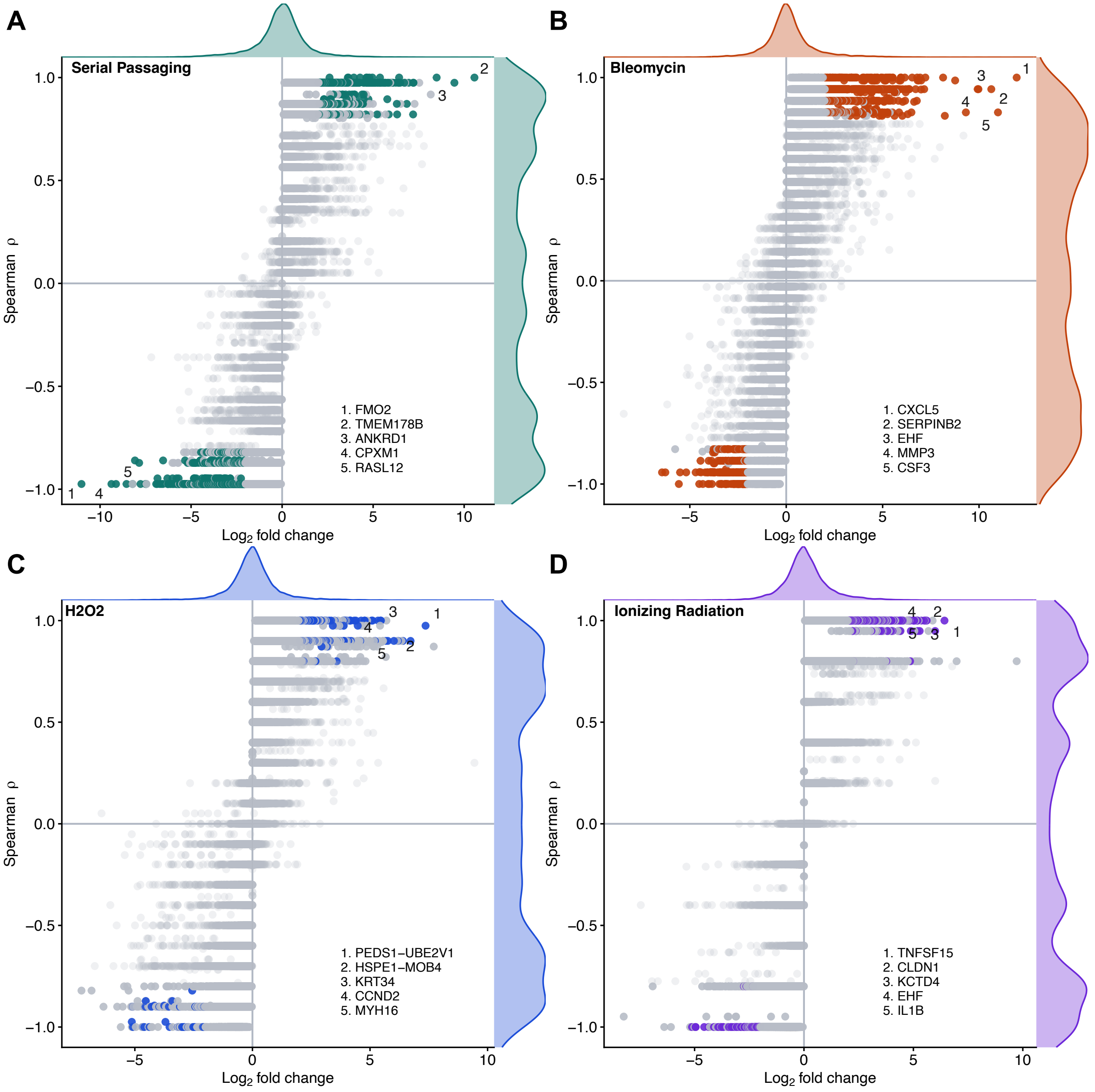


**Figure S8: Senescence induction leads to the differential expression of single genes. (A-D)** Monotonicity and fold-expression change of single genes induced by SP, BL, OX, or IR. Monotonicity was calculated using the Spearman correlation (ρ) of a linear model between $\log_2$(normalized count + 1) and days in culture or ln(dose + 1), as well as the associated *p*-value, while $\log_2$-fold change in gene expression was calculated between the highest- and lowest-PDL or dose samples. Differentially expressed genes are displayed in non-gray colors, with numerical rankings highlighting the five genes with the highest magnitude fold-change within each treatment.

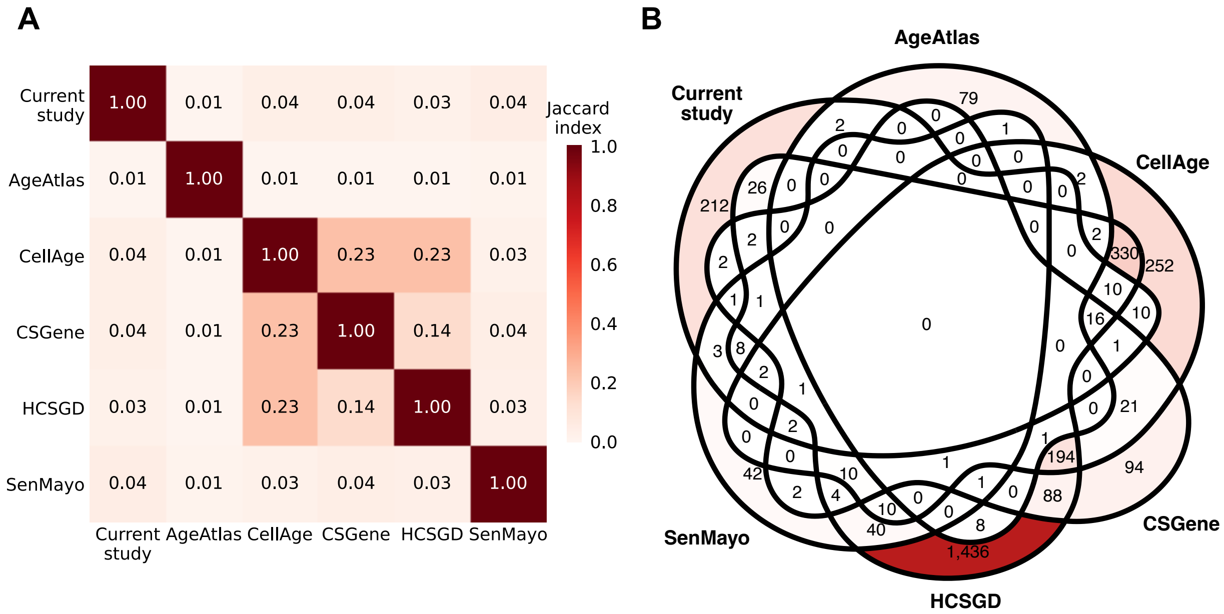


**Figure S9: Senescence-associated gene sets exhibit minimal overlap with each other. (A)** Overlap between genes identified by the current study, as well as from other AgeAtlas, CellAge, CSGene, HCSGD, and SenMayo. Genes from the current study were selected based on significant differential expression across at least two inducers. Overlap was quantified as the Jaccard index between pairs of gene lists. **(B)** Venn diagram of the overlap between senesce-associated gene sets.

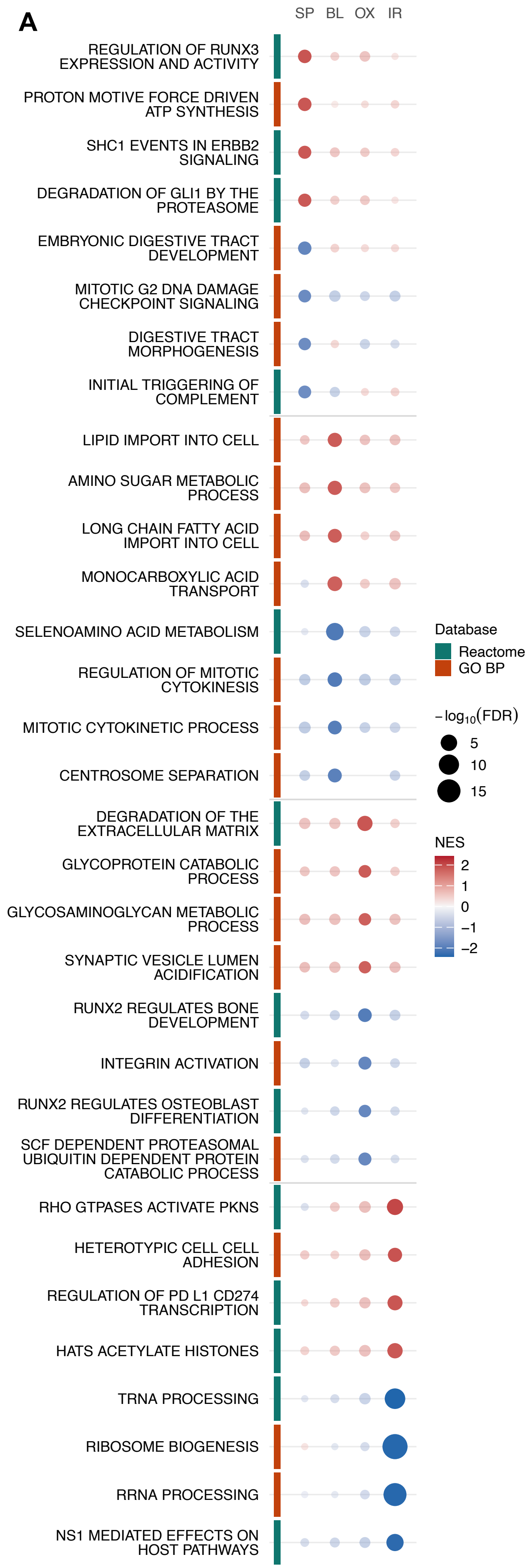


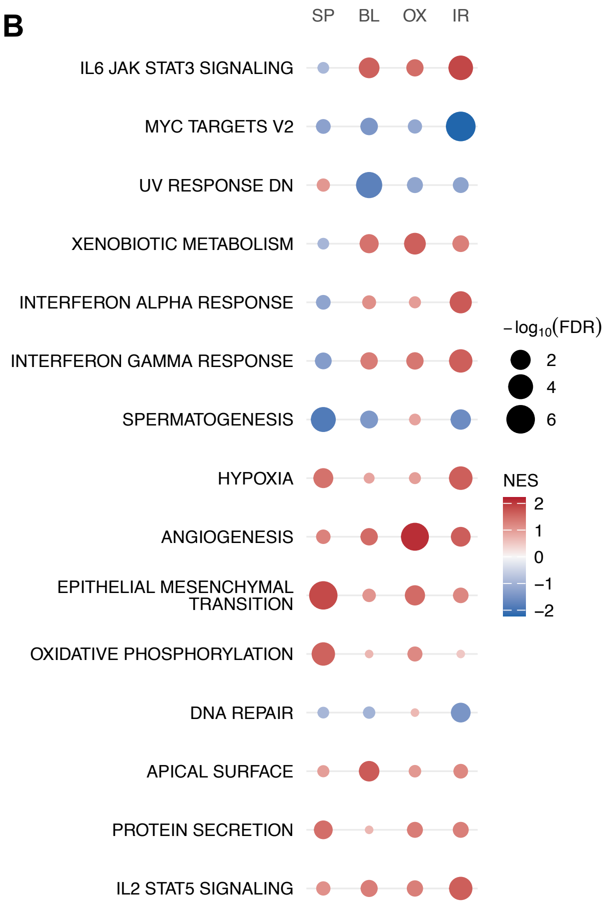


**Figure S10: Enrichment of some senescence-associated pathways varies by inducer. (A)** Top 4 positively and negatively enriched gene sets unique to each treatment. Pathways were ranked by their treatment-specific normalized enrichment scores (NESs), with treatment-specific significance thresholds of Benjamini-Hochberg-adjusted False Discovery Rate (FDR) ≤ 0.05. Gene sets are labeled with their names and databases of origin. **(B)** Enrichment of the 15 most variably enriched Hallmark gene sets across all treatments. Gene sets were ranked by the standard deviation of their treatment-specific NESs.
